# Segment Based Pattern Analysis Reveals a Persistent Regular Rhythm in the Motion Artifact Record of the MIT-BIH Noise Stress Test Database

**DOI:** 10.1101/2022.10.18.512701

**Authors:** Bruce Hopenfeld

**Affiliations:** Seekuence Solutions, New Harmony UT, USA

## Abstract

**Background:** The MIT-BIH Noise Stress Test Database (NSTDB) is a publicly available resource for testing QRS detection algorithms. Serial QRS detection algorithms applied to the NSTDB have apparently failed to detect the presence of a possible heartbeat like rhythm associated with peaks that the NSTDB classifies as noise. The failure to detect this rhythm may arise from the difficulty associated with interpreting noisy RR interval time series produced by serial QRS detection schemes.

**Algorithm Summary and Experiment:** To extract rhythm information from noisy peak time/RR interval time series, a peak space signal is created with triangular pulses centered on peaks located by a serial QRS detection algorithm such as Pan-Tompkins. The peak space signal is autocorrelated over 20 s segments and the primary (non-origin) peak in the autocorrelation signal is located. In the presence of reasonably regular sinus rhythm, this peak corresponds to a fundamental RR interval present throughout the 20 s segment. This peak time processing method was applied to the Pan-Tompkins QRS detections in the motion artifact record of the NSTDB. To compare the results to a different algorithm capable of detecting patterns at the segment level, a previously described pattern-based heartbeat detection scheme (Temporal Pattern Search, or “TEPS”) was applied in both single and multiple channel modes to the NSTDB motion artifact record.

**Results:** Both the Pan-Tompkins/autocorrelation method and TEPS detected a persistent rhythm around 1000-1050 ms in both channels throughout the entirety of the motion artifact record. The RR interval correlation between Pan-Tompkins/autocorrelation and single channel TEPS was 0.8 and 0.7 in channels 1 and 2 respectively with p values of 0.

## 1. Introduction

Serial QRS detection algorithms can produce RR interval time series with outliers that may be removed by smoothing or filtering[1]. However, when noise is very high, the RR interval time series may be so erratic that such smoothing techniques will not work. To overcome challenges associated with serial detection, neural networks have been applied to segments of ECGs to generate a sequence of QRS complexes in parallel[2]. Alternatively, a methodology known as TEPS (for Temporal Pattern Search) involves performing a combinatorial search for an optimal peak sequence within a data segment [3,4,5,6]. One challenge in benchmarking these alternatives is the relative paucity of publicly available databases that include very noisy ECGs with known QRS times.

The Physionet Noise Stress Test Database (NSTDB) is one such database[7,8]. The NSTDB includes three half hour two channel signals corresponding to three different types of noise: baseline wander, electromyogram, and electrode motion (motion artifact). Figure 1 shows a portion of the motion artifact signal. The NSTDB also includes clean ECGs from the MIT-BIH Arrhythmia Database [9] to which has been added calibrated amounts of the three noise signals. If these records are to serve as the basis for assessing the noise resistance of algorithms, it is imperative that the added noise does not include a substantial number of true QRS complexes. Otherwise, an algorithm that detects such QRS complexes would be unfairly penalized.

**Figure 1.**
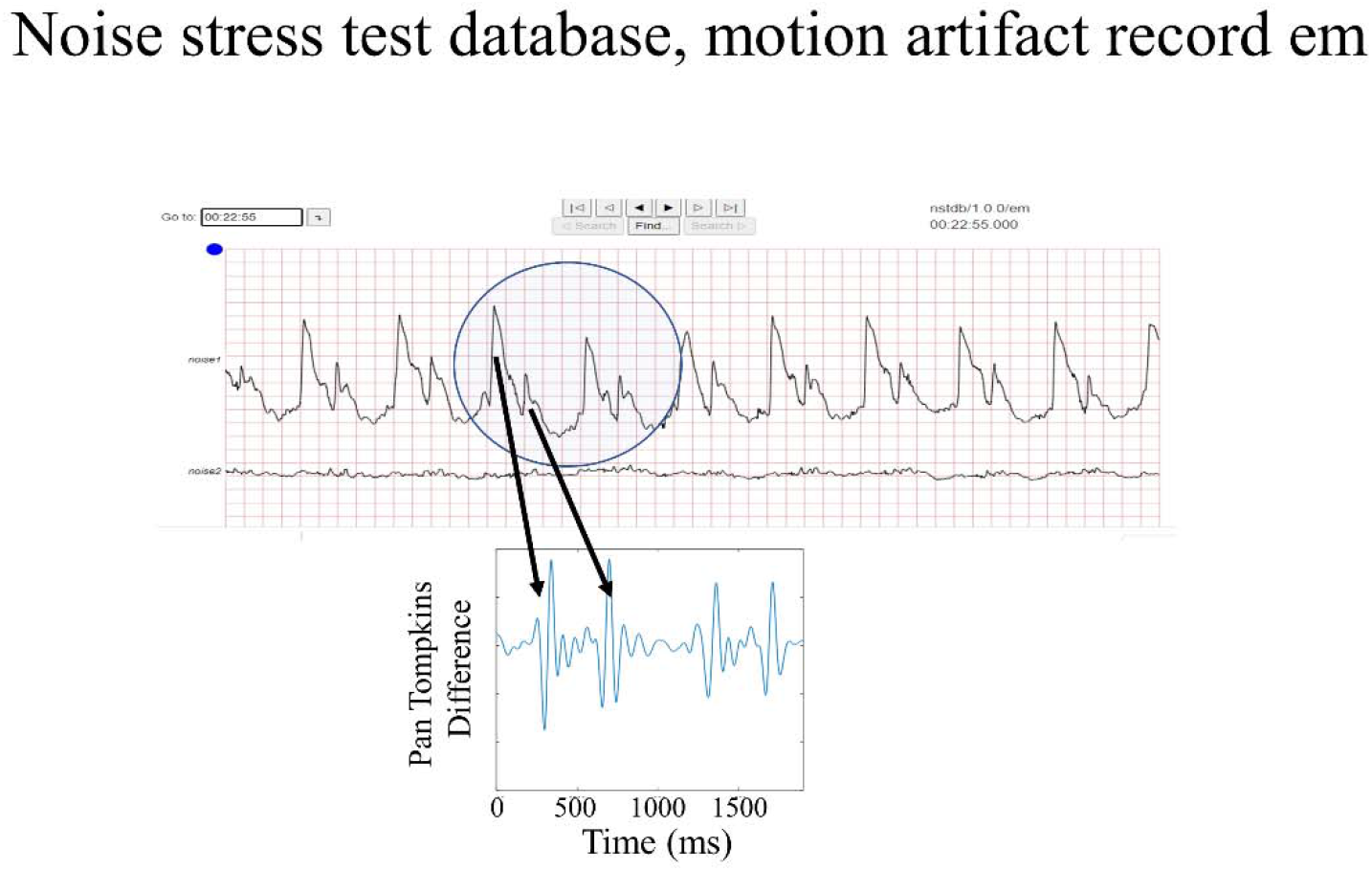
Plot of a segment from channel 1 of the NSTDB motion artifact signal as displayed by the Physionet Lightwave viewer (https://physionet.org/lightwave/). The inset shows the Pan-Tompkins five point first derivative/difference of a portion of the segment. The arrows indicate QRS and T wave like portions of the signal. (In the Lightwave screen shot, the lower amplitude channel 2 is shown below channel 1.)

Motion artifact spikes can mimic the shape and size of QRS complexes[7]. Methods for avoiding false positives for such QRS mimicking noise include: 1) tagging them as noise by correlating them to an exogenous noise-related signal (such as an accelerometer) [5], 2) determining whether such peaks temporally cohere across different ECG channels[4, 6]; and 3) relying on *a priori* aspects of heart rhythm that aren’t mimicked by sequences of random motion artifact noise [3,4,5,6]. In a single channel recording, option 2 is not available, and option 3 can be confounded if the motion artifact is rhythmic, in which case, only option 1 is viable. Thus, if a single channel contains rhythmic sequences of QRS like spikes, QRS detection that relies on option 3 will not work unless an accelerometer (or other noise related) signal is available. The NSTDB does not include accelerometer signals. Consequently, if the NSTDB motion artifact signal includes rhythmic sequences of QRS like spikes, QRS detection algorithms will struggle to avoid false positive detections. (In low/moderate noise conditions, peak shape clustering, e.g. as described in [10], along with very informative *a priori* peak shape probabilities could potentially detect sequences of true QRS complexes amongst sequences corresponding to lower probability QRS shapes.)

As will be shown in this work, the NSTDB motion artifact signal indeed contains persistent sequences of rhythmic QRS spikes, as suggested by Figure 1 and as indicated by the similar results produced by: 1) a new autocorrelation method applied to the output of a serial QRS detector.; and 2) single and multiple channel TEPS.

## 2. Algorithm/Experiment

Two different algorithms were applied to both channels of the NSTDB motion artifact signal: 1) Pan-Tompkins followed by peak space autocorrelation over 20 s segments to derive average RR intervals; and 2) TEPS.

Pan-Tompkins (from Hooman Sedghamiz, available at https://www.mathworks.com/matlabcentral/fileexchange/45840-complete-pan-tompkins-implementation-ecg-qrs-detector) is applied to 20 s long signal segments centered 10 s apart from one another. A triangular peak pulse train signal is generated from these peaks[6], with each triangle centered on a corresponding peak time. The triangles are of unit amplitude and 200 ms wide at the base, wide enough to allow reasonable RR interval changes to be captured by autocorrelation; Figure 2 shows an exemplary peak pulse train. The pulse train signal is autocorrelated, and the primary peak (along the positive x axis) in the autocorrelation signal is detected. This peak corresponds to repeated inter-peak intervals (possibly RR intervals) in the segment. An RR interval time series (10 s increments) is derived from these autocorrelation peaks, and outliers are removed. Additionally, pulse trains (for the same 20 s segment) corresponding to two different channels are cross correlated to provide a measure of peak time coherence across channels’ peaks[6].

**Figure 2.**
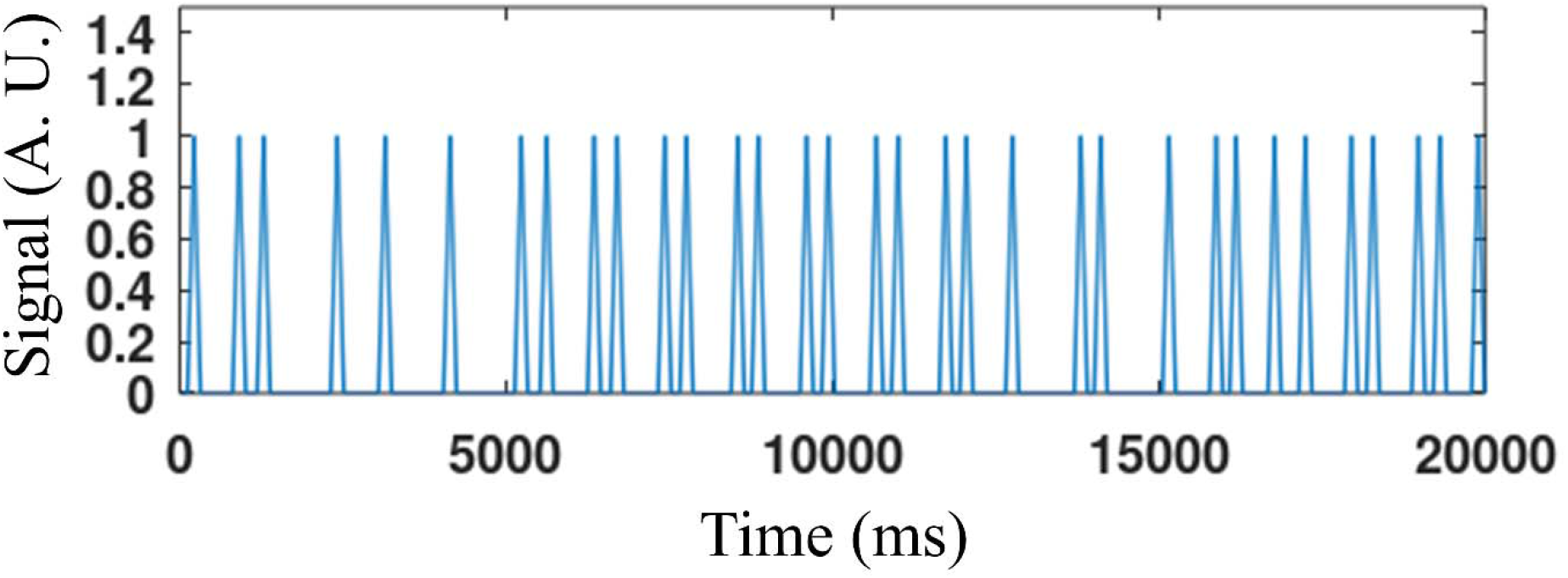
A triangular pulse train corresponding to the peaks located by the Pan-Tompkins algorithm in the first 20 s of channel 1 of the NSTDB.

Single channel TEPS was separately applied to the two NSTDB motion artifact signal channels. In addition, the two channels were processed by multi-channel TEPS, which takes advantage of peak time coherence across channels and merges all peaks across channels. Both single and multi-channel TEPS have been previously described in [4,5,6]. Briefly, according to single channel TEPS, ECG sequences are generated from candidate peaks and scored for quality according to: 1) prominence of the peaks within a sequence; 2) temporal regularity; and 3) the number of skipped beats. In multiple channel TEPS, ECG sequences are generated from a combination of peaks across channels and these sequences are further scored according to the timing coherence of peaks between channels. The temporal regularity score component is a measure that quantifies the change in between peak time intervals over a sequence.

The raw sequence score SC is set equal to:

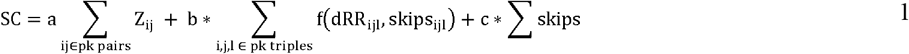

where Z is the prominence ratio for each peak pair in a sequence, dRR_ijl_ is the change in RR intervals between 3 consecutive peaks with corresponding skipped beats skips_ijl_, f() is a Gaussian with a standard deviation that depends on skips_ijl_. Additional details are included in [4].

For each segment, TEPS produces a set of peak time sequences and associated quality scores SC. In the present work, a median RR interval was computed for the highest scoring sequence for each segment, resulting in a sequence of RR intervals RRT(i) with associated scores SC(i), where “i” is a segment number index. The final RR interval time series RRU is derived by regularizing a score weighted RRT according to a second derivative operator L:

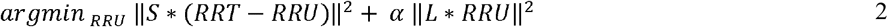

where S is a diagonal matrix corresponding to the score vector SC. The solution is:

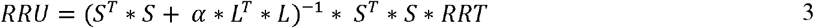

All computations were performed on a 2017 Hewlett Packard Laptop with an Intel Core i3-8130U CPU, base frequency 2.20GHz, with 8GB of RAM. For TEPS, signals are preprocessed by low pass filtering with a 5^th^ order Butterworth filter with a cutoff frequency of 45Hz, the filtered signals are downsampled to 256Hz, and the resulting signal x() is differenced according to: y(i) = x(i-12)-2*x(i-6)+x(i). The number of candidate peaks (NCP) is set equal to 30 with segment duration set at 7.5 s. Candidate Peaks must meet broad minimum and maximum peak width criteria.

## 3. Results

The upper and lower panels in Figure 3 show the RR intervals in channels 1 and 2 respectively as estimated by both Pan-Tompkins/autocorrelation and single channel TEPS. As a reference, the RR intervals estimated by multiple channel TEPS are shown in both panels. All of the methods’ RR intervals and the RR intervals between channels were in broad qualitative agreement. A rhythm around 1 Hz persists throughout the entirety of both channels. The RR interval correlation between Pan-Tompkins/autocorrelation and single channel TEPS was 0.8 and 0.7 in channels 1 and 2 respectively with p values of 0.

**Figure 3.**
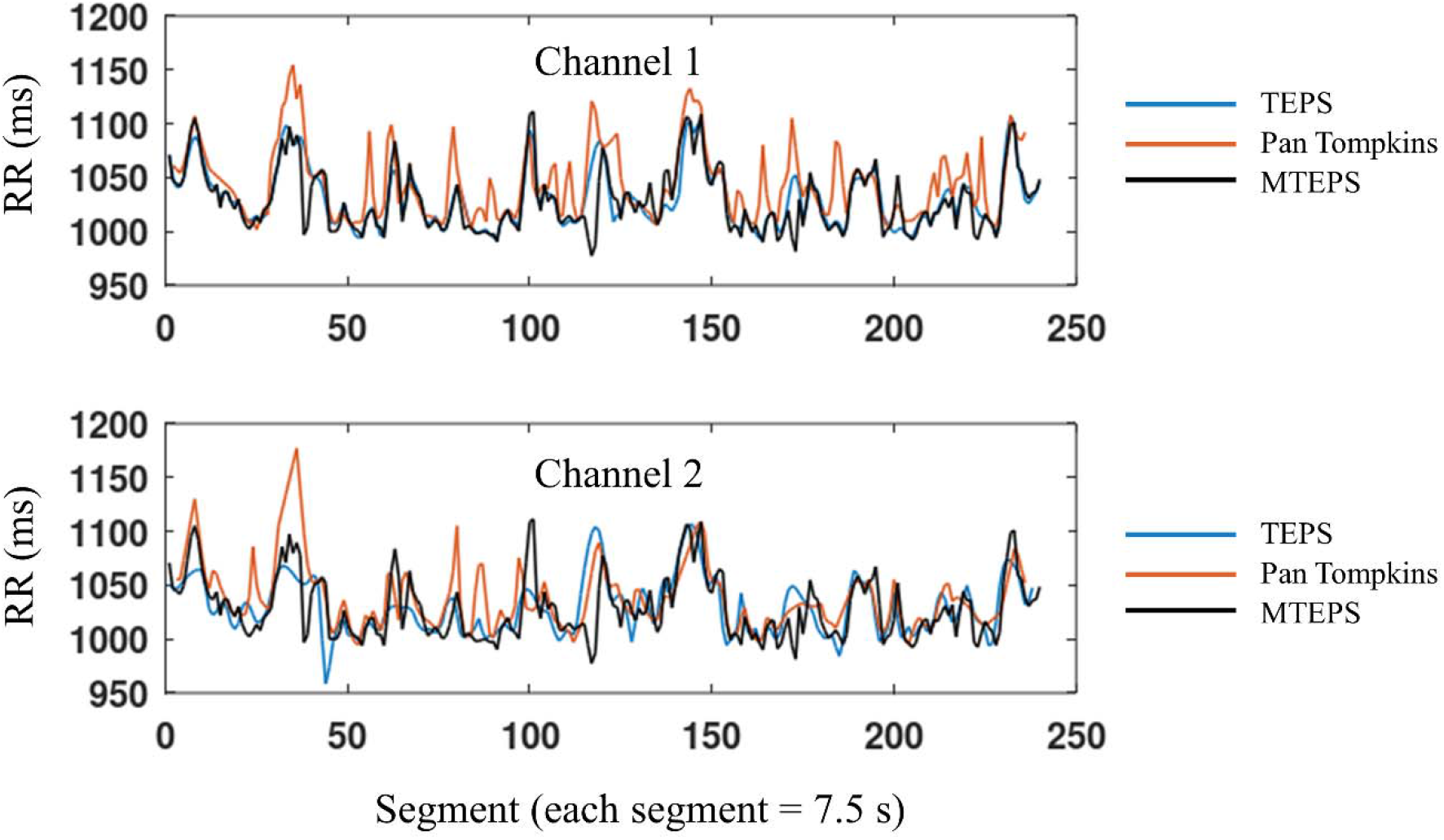
RR Interval time series associated with channel 1 (top panel) and channel 2 (bottom panel) of the NSTDB as estimated by Pan-Tompkins/autocorrelation and single channel TEPS. Both plots also include copies of the RR time series from multiple channel TEPS, which combines information from both channels.. (Autocorrelation was applied to 20 s segments and then interpolated to 7.5 s segments to match the TEPS segment duration.)

Figure 4 is a histogram of channel 1 Pan-Tompkins/autocorrelation inter-peak intervals (termed “RR intervals”) before application of autocorrelation post-processing. As shown, there are peaks in the histogram around 300 ms and 700 ms respectively. There is a smaller peak at approximately 1000 ms. Figure 5 shows a portion of the channel 1 autocorrelation for the first segment (i.e, time 0 s through 20 s). There is a peak at approximately 1000 ms and harmonics of this interval. Consistent with the Figure 4 histogram, there are subsidiary peaks at approximately320 ms and 700 ms respectively. A discrete cosine transform of the triangular peak signal produced very similar results in the corresponding frequency domain.

**Figure 4.**
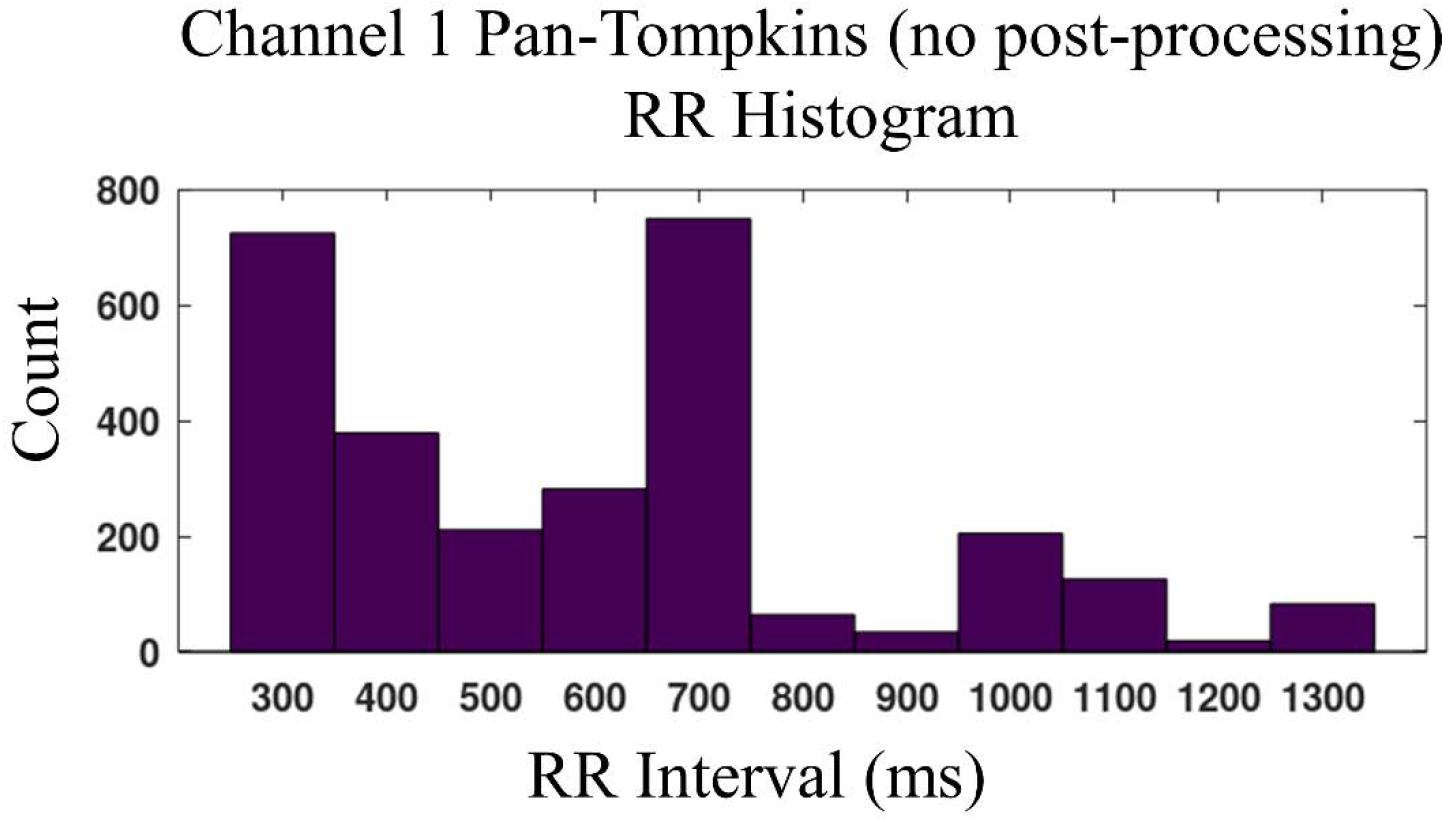
Histogram of channel 1 Pan-Tompkins without post-processing inter-peak intervals

**Figure 5.**
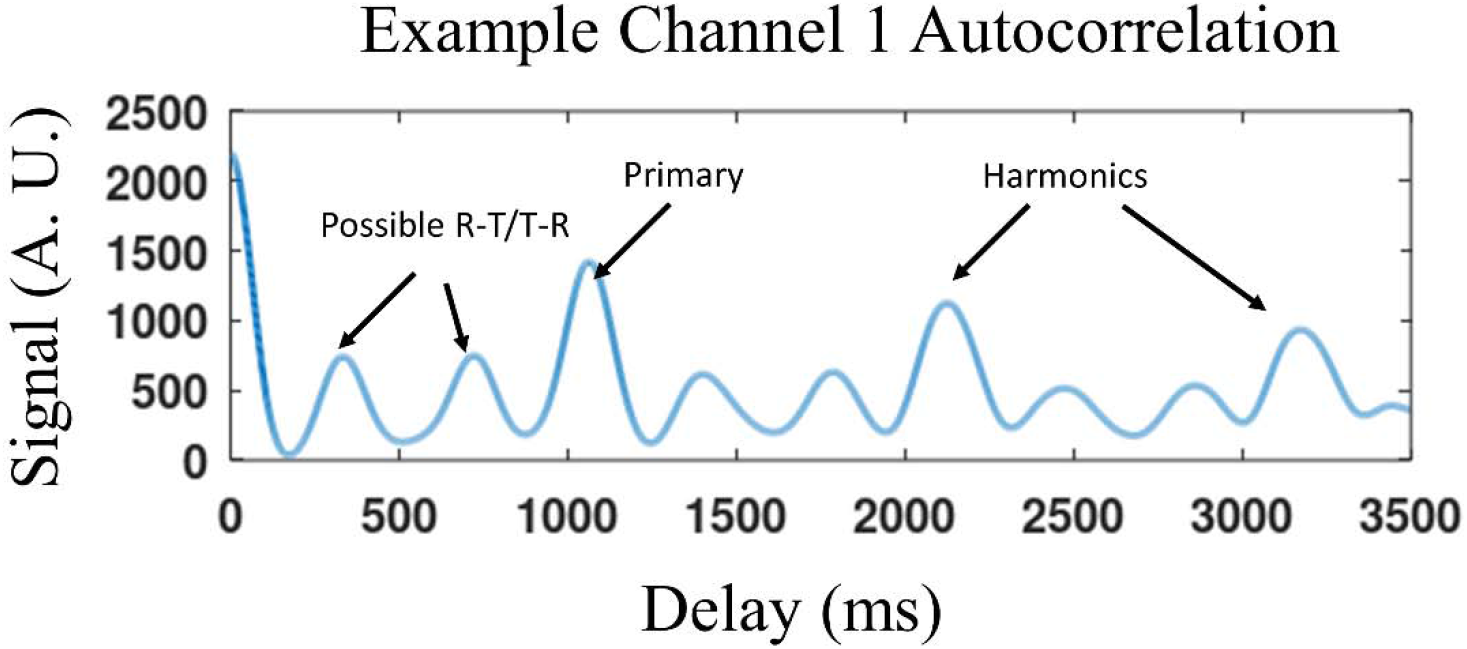
A portion of the channel 1 Pan-Tompkins/autocorrelation for the first segment (i.e, time 0 s through 20 s). The sum of the subsidiary peak times, approximately 320 ms and 700 ms, is approximately equal to the primary peak time of approximately 1020 ms.

Figure 6 shows the cross correlation maximizing offset between channels 1 and 2 over time. Outliers have been removed. There is roughly a 60-80 ms peak time offset between the channels that persists throughout the record. The cross correlation also showed a secondary peak at the 1000-1050 ms base rhythm.

**Figure 6.**
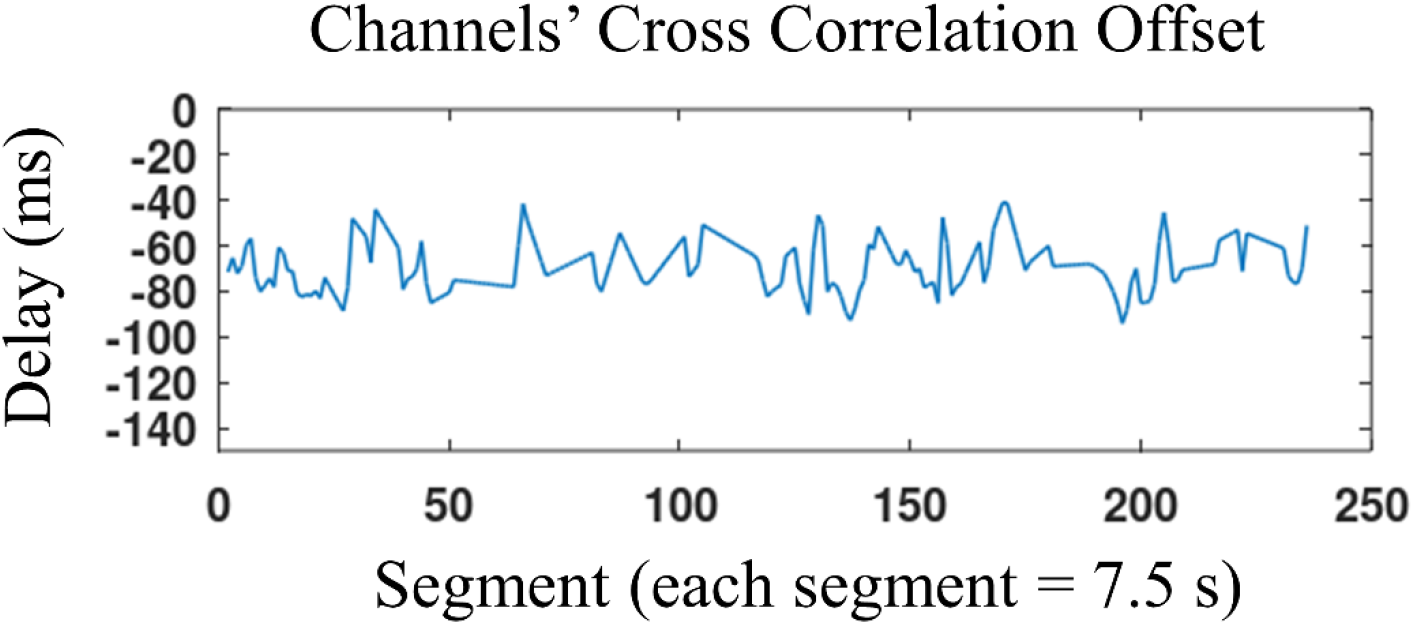
The cross correlation maximizing offset between channels 1 and 2 over time. Outliers have been removed. (Cross correlation was applied to 20 s segments and then interpolated to 7.5 s segments to match the TEPS segment duration.)

## 5. Discussion

The results strongly suggest that there is a persistent heartbeat like rhythm in both channels of the motion artifact record of the NSTDB. This rhythm is either caused by a heartbeat or an external noise source. Some aspects of the results are consistent with a heartbeat: 1) the 320ms subsidiary interval (Figure 4) is within the range of the normal adult human R to T wave interval [11]; 2) the rhythm would appear to be unusually consistent for most types of motion artifact noise; 3) portions of the signal such as those shown in Figure 1 are quite clean apart from the “noise” waveforms; 4) portions of the waveforms show very intricate patterns, such as the notches in the possible T waves shown in Figure 1; 5) the rather stable peak time offset between channels suggests a source in fixed relation to the recording electrodes. Although the QRS like waveforms are oddly shaped and correspond to unusually long QRS durations, these abnormalities could be a result of a pathological condition such as a bundle branch block[12]. In any event, any QRS detection algorithm that tags these peaks as noise will very likely be unable to detect as QRS complexes the unusual shapes associated with e.g. record 207 of the MIT-BIH Arrhythmia database. Further, in high noise conditions, true QRS complexes are often embedded within oscillatory noise, which means that waveforms such as those shown in Figure 1 cannot be discarded as noise without also eliminating true QRS complexes.

In portions of the high noise (0 dB, −6dB) records of the NSTDB, a segment based QRS detection algorithm such as TEPS may locate the 1000 ms motion artifact noise sequences, which are considered false positives. The results in this work suggest that such false positives are an improper measure of a QRS detection algorithm’s performance.

## Notes

### Competing Interest Statement

The authors have declared no competing interest.

